# Personal transcriptome variation is poorly explained by current genomic deep learning models

**DOI:** 10.1101/2023.06.30.547100

**Authors:** Connie Huang, Richard Shuai, Parth Baokar, Ryan Chung, Ruchir Rastogi, Pooja Kathail, Nilah Ioannidis

## Abstract

Genomic deep learning models can predict genome-wide epigenetic features and gene expression levels directly from DNA sequence. While current models perform well at predicting gene expression levels across genes in different cell types from the reference genome, their ability to explain expression variation between individuals due to cis-regulatory genetic variants remains largely unexplored. Here we evaluate four state-of-the-art models on paired personal genome and transcriptome data and find limited performance when explaining variation in expression across individuals.

## Main

With rapid advances in deep learning and growing datasets for training, there has been recent success in predicting gene expression levels [1–4], 3D genome folding [5, 6], and epigenetic features [7–10] such as transcription factor binding, histone modifications, and chromatin accessibility directly from the reference genome sequence. These genomic deep learning models are trained using genome-wide data from a variety of cell types and cellular contexts and have been shown to learn biologically relevant regulatory motifs within the input DNA sequence [8, 9]. Current sequence-to-expression models can explain variation in expression across different genes in the genome based on the reference genome sequence surrounding the start site of each gene. However, the application of such models to sequences from personal genomes to explain variation in gene expression across individuals (Fig. 1a) has been largely unexplored. Here we evaluate four state-of-the-art models—Enformer [4], Basenji2 [11], ExPecto [2], and Xpresso [3]—on paired whole genome sequencing and RNA-sequencing data (*n* = 421) from the Geuvadis consortium [12] and show that model performance is limited when explaining gene expression variation across individuals. When the models do pick up on regulatory variation, for a limited set of genes, they often fail to capture the correct direction of effect of such variation on expression. Together with the recent findings of Sasse, Ng, Spiro *et al*. [13], our work highlights shortcomings of current deep learning models of gene expression when applied to personal genome interpretation.

**Fig. 1:**
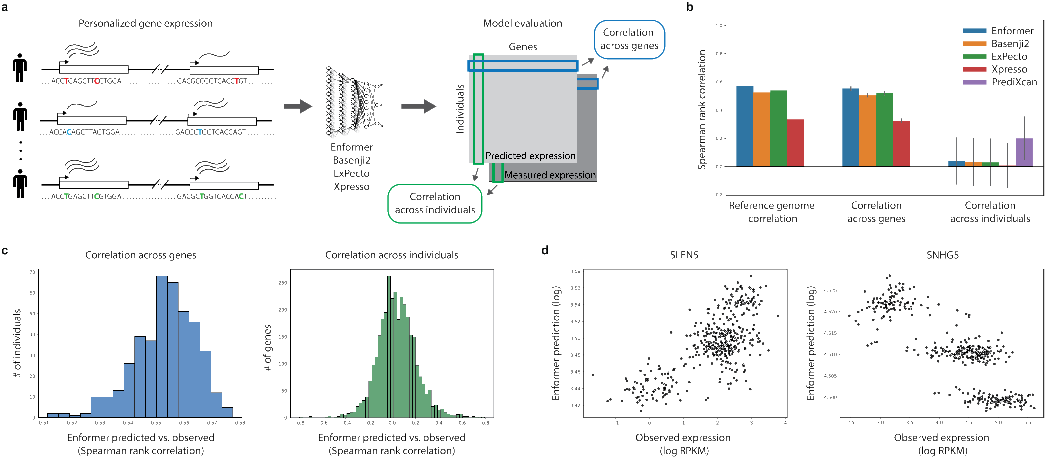
Cross-gene vs. cross-individual gene expression prediction. (a) Overview of our approach. (b) Performance of all tested models on reference sequence prediction, cross-gene prediction averaged across individuals, and cross-individual prediction averaged across genes. Error bars represent the standard deviation over all individuals or genes, respectively. (c) Enformer cross-gene performance for each individual (left histogram, distribution over individuals) and Enformer cross-individual performance for each gene (right histogram, distribution over genes). Histograms for the other tested models are shown in Fig. S2 and S3. (d) Example genes with strong positive correlation (left) and strong negative correlation (right) for Enformer.

To test these existing sequence-to-expression models on personal genome variation, we use RNA-sequencing data from the Geuvadis consortium, measured on lymphoblastoid cell lines (LCLs) and paired with whole genome sequencing (WGS) data from 421 individuals in the 1000 Genomes Project [14]. We focus on the 3,259 genes for which the Geuvadis analysis of expression quantitative loci (eQTLs) identified at least one statistically significant genetic association where genotype of a cis variant is predictive of gene expression variation across individuals. We construct personal input sequences for each individual by inserting their single nucleotide variants (SNVs) into the reference sequence around each gene transcription start site (TSS). We then compute gene expression predictions for each individual, as well as for the reference genome sequence, using all four models (see Methods for details). For each model, we use the output expression prediction track corresponding to the cell type most similar to the LCLs used for the Geuvadis measurements. To ensure that the chosen model outputs are indeed relevant for prediction of gene expression in LCLs, for each gene we compare the model prediction using the reference sequence to its median expression level in the Geuvadis dataset (Fig. 1b and S1). We find Spearman rank correlations between predicted and observed expression levels of 0.57 for Enformer, 0.52 for Basenji2, 0.54 for ExPecto, and 0.33 for Xpresso, indicating that these models explain a significant fraction of expression variation across genes in LCLs, similar to previous reports.

For each model, we then compute two additional metrics using the personalized sequences as input. First, for each individual, we calculate a cross-gene correlation that compares the predicted expression levels of the aforementioned 3,259 genes using that individual’s personal input sequence to the observed expression levels of those genes in the same individual. Similarly, for each gene, we compute a cross-individual correlation that compares the predicted expression levels in all 421 individuals to their observed expression levels (see Fig. 1a for a visual comparison of the two metrics). We find that the cross-gene correlation for each individual is similar to the reference genome performance of the corresponding model (Fig. 1b,c and S2), with average Spearman correlations of 0.55 for Enformer, 0.51 for Basenji2, 0.52 for ExPecto, and 0.32 for Xpresso. However, when we instead compute the correlation across individuals for each gene, we find that the distribution of cross-individual correlations is centered close to zero for all models (Fig. 1b,c and S3, S4), indicating that all models struggle to explain variation in expression across individuals. This result suggests that current state-of-the-art sequence-to-expression models do not correctly predict the effects of many single nucleotide variants on gene expression. We also try ensembling the predictions from the four models and find that performance is only slightly improved by averaging predictions across models (Fig. S5).

In comparison, regularized linear regression models trained separately for each gene using nearby variant dosages as predictors (the approach used by PrediXcan [15]) explain much more cross-individual variation, even when restricted to the same input context (197kb) as Enformer (Fig. 1b and S3). Since such PrediXcan-style models do not attempt to learn generalizable sequence features that can be applied to new sequences, variants, and genes outside of the training set, we include these models not as a competing approach, but rather as a minimum baseline for the genetic contribution to expression that should be possible to learn for each gene in the dataset. The higher performance of these PrediXcan-style models indicates effects of common cis-regulatory variants that are not captured by current deep learning models.

We also find that although the mean cross-individual correlation is close to zero for all models, there are tails of strongly positively correlated and strongly negatively correlated genes for each model (Fig. 1c and S3). Example genes with strong positive correlation and strong negative correlation are shown for Enformer in Fig. 1d. When comparing predictions for such genes across all four models, we find that the models often disagree with one another on the direction of correlation (Fig. 2a,b and S6, S7). This result suggests that the incorrectly predicted direction of genetic effect for the negatively correlated genes for any given model is not due to an inherent difficulty in modeling those particular genes or their corresponding variants, but rather to noise in the effects attributed to variants by these types of models. Importantly, we find that the four tested models are more consistent with one another in the magnitude of their correlation to observed expression of a given gene than in the direction of that correlation (Fig. 2b and S6), suggesting that they agree on identifying causal regulatory variants more than they agree on the direction of effect of such variants on expression.

**Fig. 2:**
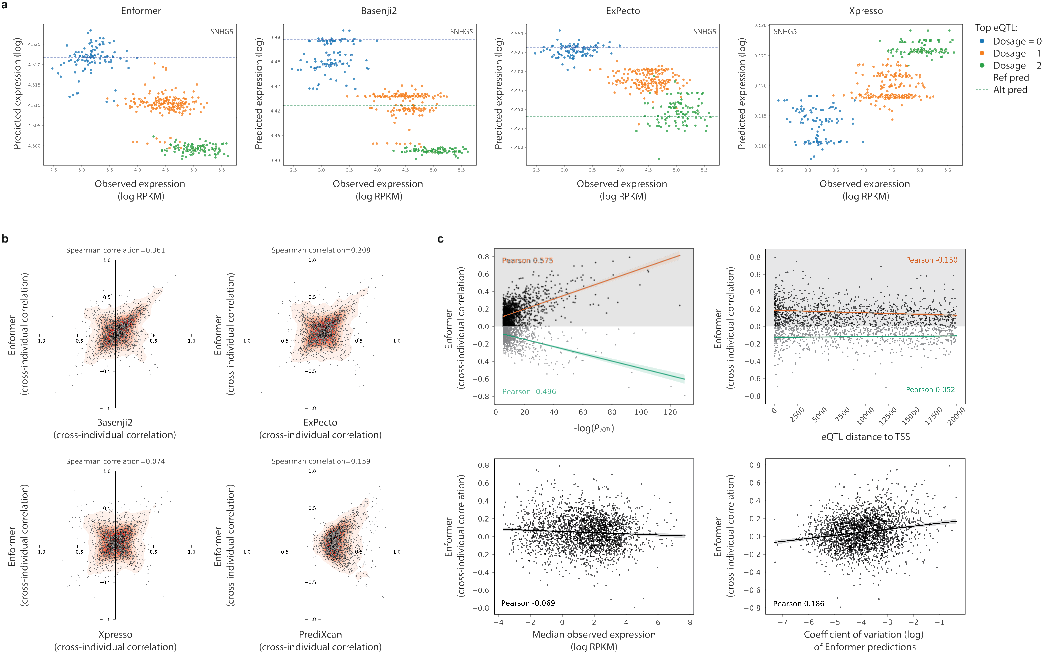
Models often disagree on predicted direction of effect of cisregulatory variation. (a) Predictions from all four deep learning models on an example gene, *SNHG5*, that has strong negative cross-individual correlations for Enformer, Basenji2, and ExPecto, and positive cross-individual correlation for Xpresso. Points are colored by each individual’s dosage of the most significant eQTL for this gene. Dashed lines represent the predicted expression levels of the reference and alternate alleles of the most significant eQTL. (b) Comparison of cross-individual Spearman rank correlations for Enformer vs. other models. Note the increased density of genes along the *y* = *x* and *y* = *−x* axes. (c) Cross-individual correlations for Enformer compared to the *p*-value of the most significant eQTL in each gene (top left), the distance to the TSS for that eQTL (top right), the median observed expression of the gene (bottom left), and the coefficient of variation of the predicted expression levels of the gene (bottom right). Note that negative correlations are observed even for genes with strong eQTLs.

We next explore whether predicted directions of genetic effect on expression tend to be more accurate for certain types of genes. First, we test whether genes with strong genetic associations in the Geuvadis eQTL analysis are more likely to have correctly predicted directions of genetic effect by comparing the cross-individual correlation for each gene to the *p*-value (Fig. 2c and S8), effect size (Fig. S9), and minor allele frequency (Fig. S10) of the most significant eQTL within 20kb of the TSS. We find that genes with strong eQTLs tend to have larger magnitude cross-individual correlations for all models; however, these genes are not more likely to have positive rather than negative cross-individual correlations, indicating that the models often predict incorrect directions of effect even for genes with strong genetic effects on expression. We find a small trend towards larger cross-individual correlations for genes with smaller distance between the most significant eQTL and the TSS (Fig. 2c and S11), which aligns with previous findings that current sequence-to-expression models capture gene expression determinants in promoters more accurately than distal enhancers [16]. However, we note that genes with proximal eQTLs still frequently have strong negative cross-individual correlations, suggesting that modeling distal regulatory effects and predicting regulatory effect direction are two important, but orthogonal, areas for future modeling improvements. Lastly, we find only small trends when comparing model performance to the median observed expression level of a gene (Fig. 2c and S12) and to the variation in predicted expression levels across individuals (Fig. 2c and S13).

In conclusion, we analyze the performance of four state-of-the-art sequence-to-expression deep learning models—Enformer, Basenji2, ExPecto, and Xpresso—on personalized gene expression prediction, and find that these models consistently under-perform when predicting differences in expression for a given gene across individuals based on inter-individual variation in the input DNA sequence. We also find genes with strong negative correlations between predicted and observed expression levels, for which the models have likely identified causal regulatory variant(s) but incorrectly predicted their direction of effect. Previous evaluations of variant effect prediction with sequence-to-expression deep learning models have focused on individual variant effects, as measured by eQTL studies, or massively parallel reporter assays (MPRAs). However, MPRAs lack the complex genomic and chromatin context of endogenous gene expression, and it is difficult to identify the causal variants in eQTL studies, even with current fine-mapping approaches, resulting in effect size estimates that are not biologically meaningful for variants in linkage disequilibrium with a causal variant. By using personal genome sequences to evaluate model performance, our input sequences include all variants surrounding the TSS for each individual and thus avoid the issue of causal variant identification. Our conclusions about directionality are in line with previous tests on eQTLs [2, 4], which showed low performance on predicting the direction of effect on expression for individual variants, especially for distal eQTLs. Following [4], we confirm this finding for Enformer for fine-mapped GTEx eQTLs in LCLs (Fig. S14).

Finally, our cross-model analysis reveals that models often strongly disagree with one another on the predicted direction of genetic effects on expression and, intriguingly, that agreement between models is greater for the magnitude of cross-individual correlation than the direction of that correlation. This result further supports the conclusion that current genomic deep learning models recognize the presence of important regulatory variation in an input sequence but struggle with understanding the direction of effect of such variation. To diagnose the reasons for these errors, it will be valuable to assess whether model predictions of variant effects on other epigenetic tracks (e.g. transcription factor binding and chromatin accessibility) are more accurate than for gene expression. For example, these models may have correctly learned variant effects on more proximal phenotypes, such as individual regulatory elements, but struggle to map effects of those elements to corresponding changes in gene expression; alternatively, the models may struggle with direction of variant effects even on proximal phenotypes such as the binding of individual transcription factors. Preliminary analysis suggests that these models still have room for improvement in predicting the direction of effect of chromatin accessibility quantitative trait loci (Fig. S15). Further work to distinguish between these possibilities will help prioritize future modeling improvements to focus on understanding high-level regulatory grammar, e.g. through hierarchical models of gene expression, or to focus on more accurately learning local variant effects, e.g. by increasing sequence diversity during model training.

## Methods

### Gene expression dataset

The data used to evaluate gene expression predictions for personal genome sequences were obtained from the Geuvadis consortium [12], which includes paired gene expression and whole genome sequencing data from individuals in the 1000 Genomes Project [14]. The E-GEUV-1 release includes RNA-sequencing data from lymphoblastoid cell lines (LCLs) from a total of 465 samples. After excluding samples with unphased imputed genotypes, there were 421 Geuvadis individuals with phased whole genome sequencing data. These samples originated from five populations with ancestry in Europe and Africa. We also obtained results from the Geuvadis cis-eQTL analysis performed in the European individuals, which considered variants with MAF *>* 5% located within 1MB of a gene TSS. Except where otherwise noted, our results are shown for all 3,259 genes that had a significant eQTL in the Geuvadis EUR cis-eQTL analysis.

### Comparison of deep learning models for gene expression prediction

We test four state-of-the-art deep learning models that make gene expression predictions for an input DNA sequence. These models consider different sequence contexts, or receptive fields, when making predictions; in particular, Enformer has the widest receptive field (98.3kb upstream and 98.3kb downstream of the gene TSS), followed by Basenji2 (27.5kb upstream and 27.5kb downstream), ExPecto (20kb upstream and 20kb downstream), and Xpresso (7kb upstream and 3.5kb downstream). All models include standard convolutional layers, with additional dilated convolutional layers in Basenji2 and transformer layers in Enformer. The models also use different sources of gene expression data during training; in particular, Basenji2 and Enformer are trained using genome-wide CAGE measurements, while ExPecto and Xpresso are trained using RNA-sequencing data. Basenji2 and Enformer use multi-task learning to make gene expression predictions along with many other epigenetic track predictions in a variety of cell types, while Xpresso predicts gene expression alone. ExPecto uses a hierarchical model, making predictions of epigenetic tracks along the input sequence and then adding a linear transformation on top of those outputs to predict expression.

### Constructing personalized input sequences

For each gene, the ENSEMBL gene ID, TSS position, strand, and chromosome were obtained from Geuvadis. We used hg19 as the reference genome for creating personalized sequences, to match the Geuvadis dataset. For ExPecto, Basenji2, and Enformer, whose receptive fields are symmetric about the TSS, and for genes located on the positive strand for Xpresso, we directly computed personalized sequences around the TSS using bcftools consensus [17]. Since Xpresso uses an asymmetric input sequence, for genes located on the negative strand we extracted the reference sequence 3.5kb before the TSS to 7kb after the TSS using samtools, applied bcftools consensus, and then took the reverse complement. We considered only single nucleotide variants (SNVs) and did not include indels when creating the personalized input sequences. We then predicted gene expression levels as described below for the two personalized haplotypes for each individual and averaged the predictions from both haplotypes.

### Basenji2 predictions

Basenji2 takes input sequences of 131kb with an effective receptive field of 55kb for each prediction. The model outputs predictions in 128-bp bins for 5,313 epigenetic and transcriptional tracks from the ENCODE, Roadmap Epigenomics, and FANTOM consortiums. We used Basenji2 predictions for the GM12878 lymphoblastoid cell line CAGE track, as it is the most relevant cell type for the Geuvadis expression data. For a given input sequence centered at a gene TSS, we averaged predictions from the forward and reverse complement sequence as well as minor sequence shifts to the left and right (1 nucleotide in each direction). To compute the final expression prediction for a gene, we averaged the predicted CAGE signal over the 128-bp bin containing the TSS, the 5 bins upstream of the TSS, and the 5 bins downstream of the TSS.

### Enformer predictions

Enformer replaces the dilated convolutions of Basenji2 with a self-attention mechanism, which facilitates the learning of long-range dependencies. Enformer has a receptive field of 196.6kb and outputs predictions in 128-bp bins for the same 5,313 tracks as Basenji2. As above, we used Enformer predictions for CAGE measurements performed on the GM12878 lymphoblastoid cell line. While the Enformer authors averaged predictions within a 3-bin window around each gene TSS, we found that averaging over a 10-bin window led to better performance on the Geuvadis dataset.

### ExPecto predictions

ExPecto predicts gene expression by first using a convolutional neural network (Beluga, an updated version of DeepSEA [7]) to predict chromatin features within a 40kb region around each gene TSS. Specifically, Beluga outputs predictions in 200-bp bins for 2,002 epigenetic tracks from the ENCODE and Roadmap consortiums. To predict expression for a given gene, Beluga is first used to predict chromatin features for 200 bins centered around the TSS, averaging predictions over the input sequence and its reverse complement. The resulting predictions are spatially transformed with a set of basis functions and used as input features for an L2-regularized linear regression model to predict expression for the given input sequence. For our ExPecto predictions, we used a publicly available ExPecto model trained on EBV-transformed lymphocytes from GTEx, which we chose as the most relevant cell type to compare with the Geuvadis expression data.

### Xpresso predictions

Xpresso consists of two convolutional blocks and two fully connected layers trained on normalized RNA-seq data across 56 tissues and cell lines from the Roadmap Epigenomics Consortium. The optimal input sequence for Xpresso was found to be a 10.5kb region asymmetrically centered around the TSS. The model also includes mRNA half-life estimates as features to capture the impact of post-transcriptional regulation. For our analysis, we used predictions from the pretrained LCL-specific Xpresso model as the most relevant to the Geuvadis dataset.

### Elastic net (PrediXcan-style) gene expression model

For comparison with the sequence-to-expression deep learning models above, we also trained a cross-validated, elastic net regression model for each gene to predict Geuvadis expression measurements from common variants (minor allele frequency *≥* 0.05) within 98.3kb of the TSS to match Enformer’s receptive field. We set the elastic net mixing parameter to 0.5 and found the best regularization penalty in a 10-fold cross-validation scheme using scikit-learn’s `ElasticNetCV` [18]. We obtained individual gene expression predictions from the output of ten holdout validation splits, such that the model makes predictions on individuals not in the training set.

### QTL effect direction classification

We obtained GTEx v8 eQTLs fine-mapped using the SuSiE method [19, 20] from the Supplementary Data in Avsec *et al*. [4]. Using these data, we evaluated Enformer on its ability to predict the direction of eQTL effect on expression. We used fine-mapped eQTLs identified in EBV-transformed lymphocytes as the GTEx cell type that most closely matched the Geuvadis expression data. Using a similar method as described in Avsec *et al*. [4], we focused on variants with a posterior inclusion probability (PIP) in a credible causal set of greater than 0.9, and removed variants that affect gene expression in opposite directions for different cis-genes. We used Enformer predictions from the GM12878 CAGE track, as the closest match to EBV-transformed lymphocytes. We computed expression direction accuracy over 100 bootstrap samples from the full set of variants.

We also obtained chromatin accessibility QTL (caQTL) variants from the Tehranchi *et al*. [21] analysis of ATAC-seq data from LCLs from individuals in ten populations: four African, four European, one African-American, and one Han Chinese. Using these data, we evaluated Enformer on its ability to predict the direction of caQTL effect on accessibility. To generate accessibility predictions, we summed Enformer’s predictions from the three 128-bp bins closest to the variant for the DNase:GM12878 track (track number 69). For computational efficiency, we randomly sampled 10,000 caQTLs from the Tehranchi *et al*. dataset, and then computed effect direction accuracy over 100 bootstrap samples of those 10,000 variants.

## Data availability

The Geuvadis gene expression data and whole genome sequencing data used in this study are publicly available at https://www.ebi.ac.uk/biostudies/arrayexpress/studies/E-GEUV-1.

## Code availability

All model predictions on Geuvadis individuals and scripts to generate personalized sequences, get model predictions, and plot figures are available at https://github.com/ni-lab/personalized-expression-benchmark.

## Acknowledgements

We thank Gabriel Loeb, David Kelley, and members of the Ioannidis lab for helpful discussions. This work was partially supported by the U.S. National Institutes of Health grant R00HG009677, an Okawa Foundation Research Grant, and a grant from the UC Noyce Initiative for Computational Transformation. N.M.I. is a Chan Zuckerberg Biohub – San Francisco Investigator.

## Supplementary Figures

**Fig. S1:**
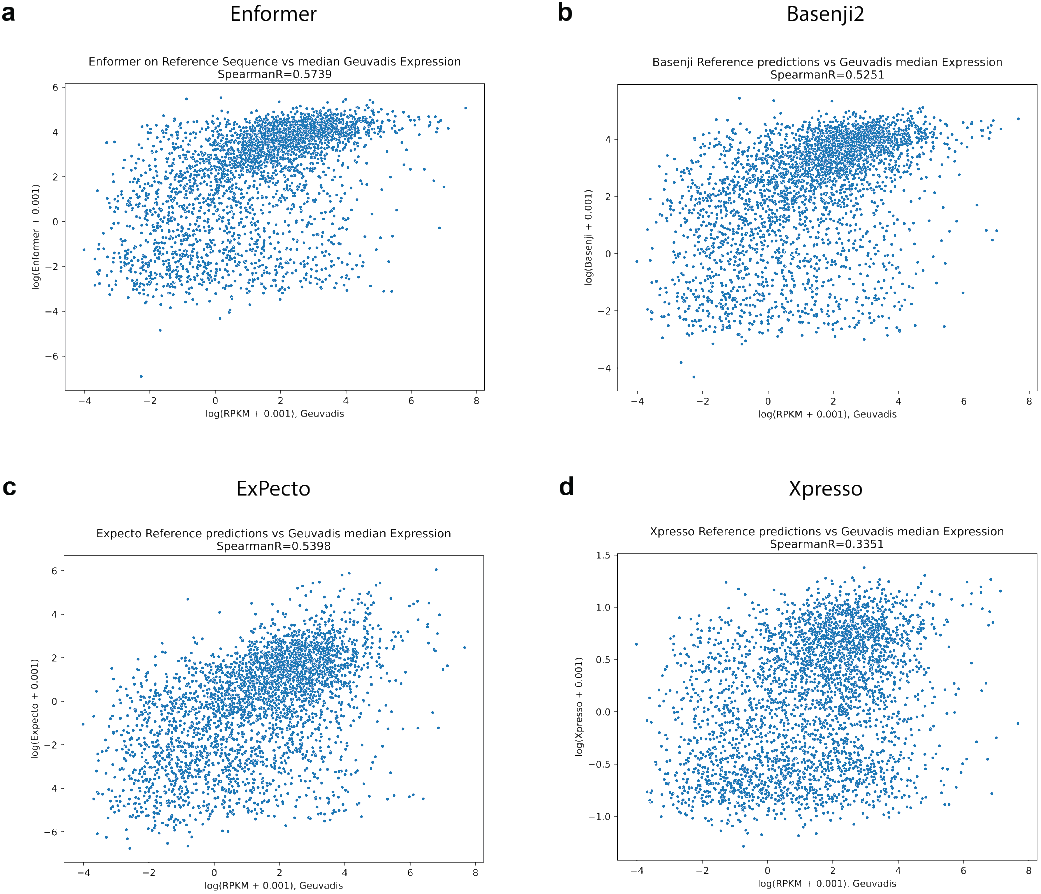
Performance of all tested models on reference sequence prediction. Median Geuvadis gene expression (log transformed) versus gene expression predictions (log transformed) obtained by inputting the reference genome sequence to (a) Enformer, (b) Basenji2, (c) ExPecto, and (d) Xpresso. For each model, gene expression predictions were used from the most relevant cell type to the Geuvadis expression data. Measurements and predictions for the 3,259 genes with at least one signficant eQTL in the Geuvadis analysis are displayed.

**Fig. S2:**
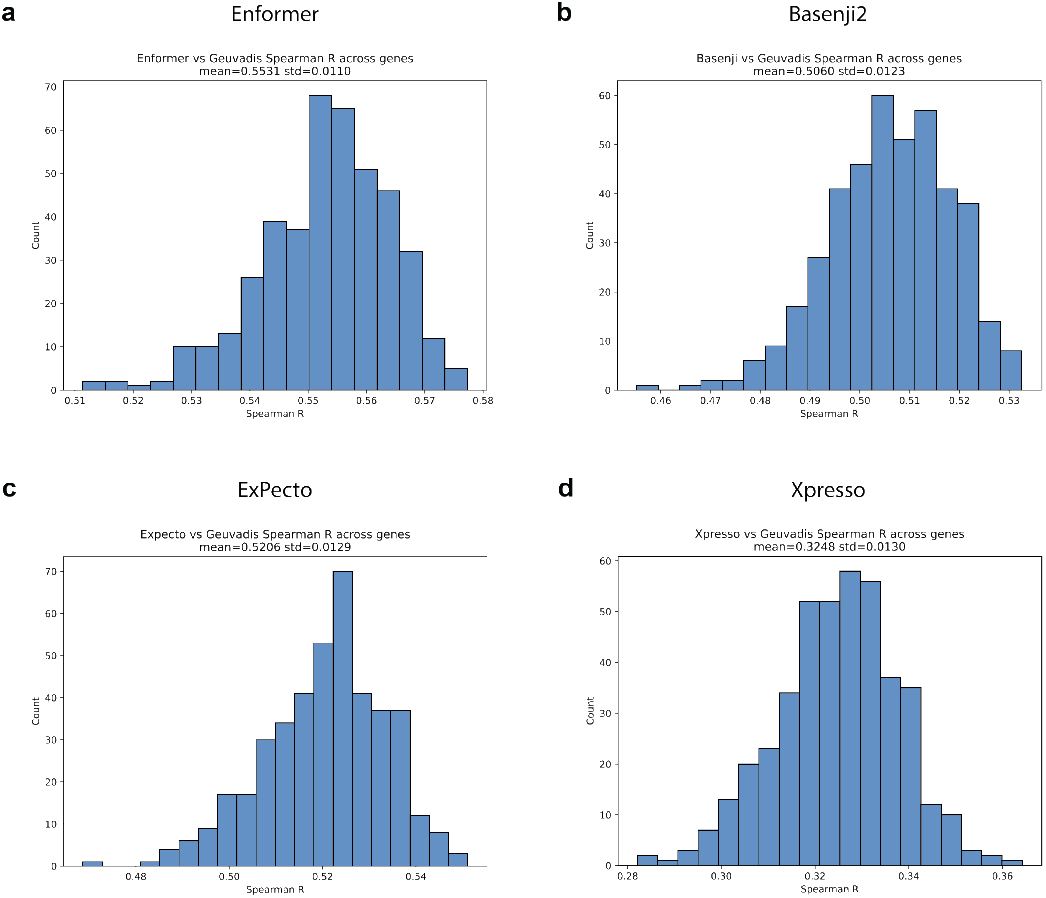
Performance of all tested models on cross-gene prediction. Cross-gene performance for each individual for (a) Enformer, (b) Basenji2, (c) ExPecto, and (d) Xpresso. For a given individual, cross-gene performance is defined as the correlation between their measured gene expression levels and gene expression predictions obtained using their personalized genome sequences. Correlations were computed across the 3,259 genes with at least one signficant eQTL in the Geuvadis analysis. Each histogram displays the distribution in cross-gene performance over individuals.

**Fig. S3:**
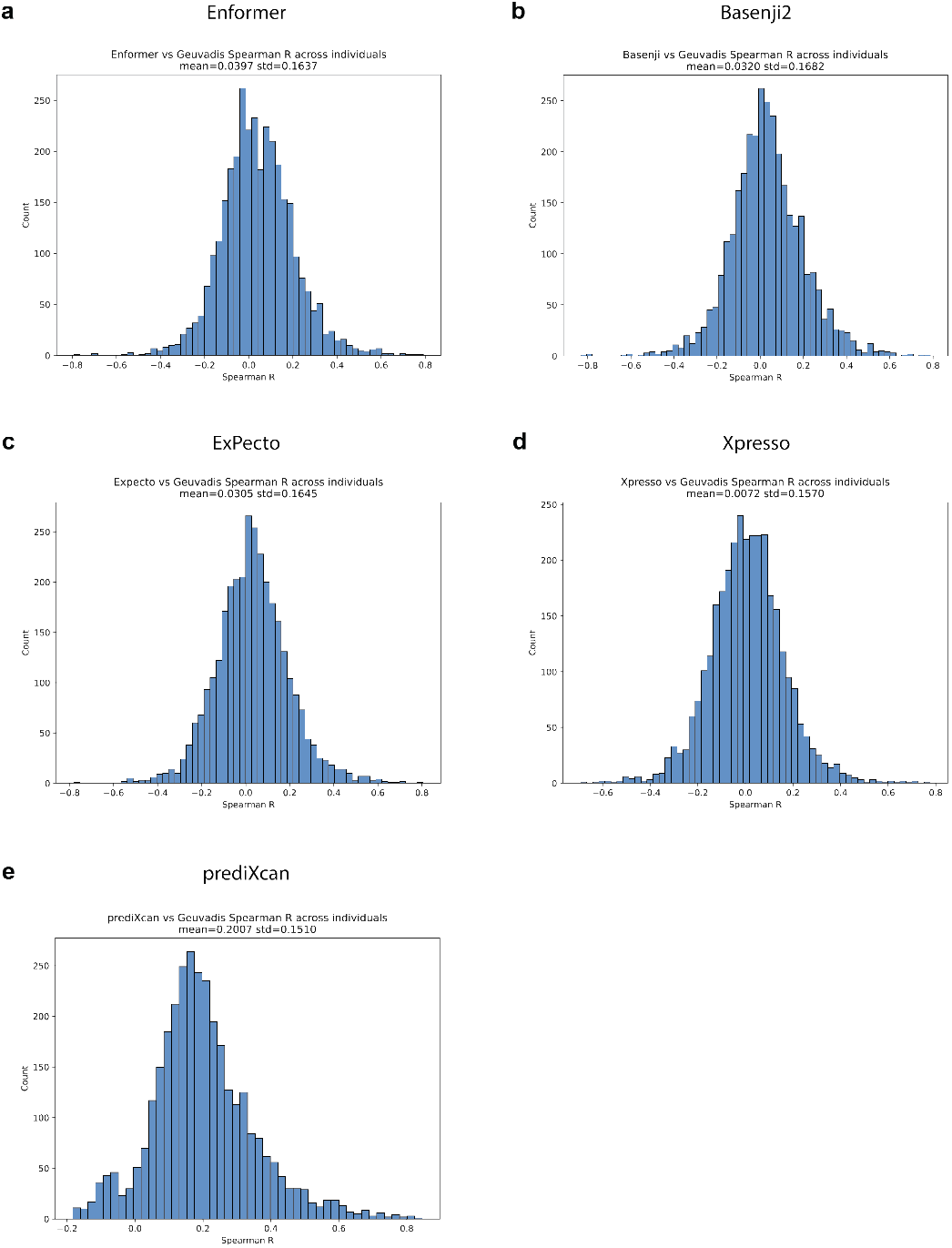
Performance of all tested models on cross-individual prediction. Cross-individual performance for each gene for (a) Enformer, (b) Basenji2, (c) ExPecto, (d) Xpresso, and (e) PrediXcan. For a given gene, cross-individual performance is defined as the correlation between measured gene expression levels in all 421 individuals and corresponding gene expression predictions obtained using each individual’s personalized genome sequence. Each histogram displays the distribution in cross-individual performance for the 3,259 genes with at least one signficant eQTL in the Geuvadis analysis.

**Fig. S4:**
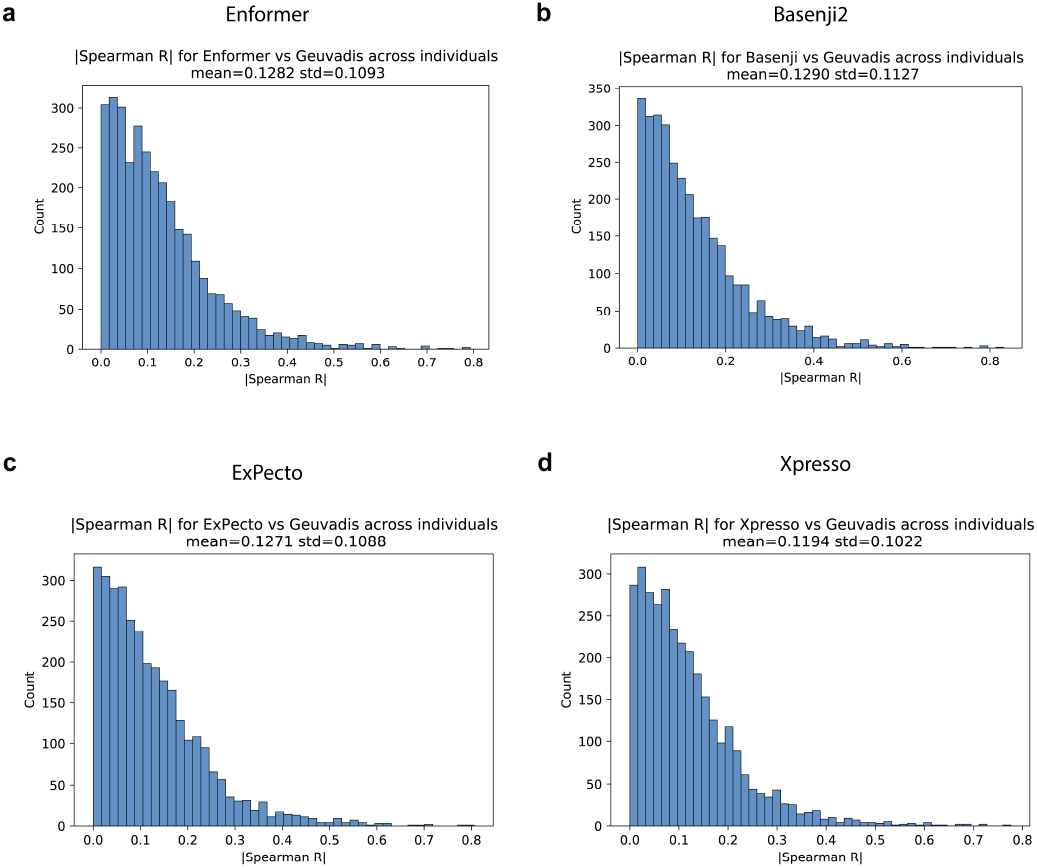
Absolute value of cross-individual correlations for all models. Absolute value of cross-individual performance for each gene for (a) Enformer, (b) Basenji2, (c) ExPecto, and (d) Xpresso. Each histogram displays the distribution in cross-individual performance for the 3,259 genes with at least one signficant eQTL in the Geuvadis analysis.

**Fig. S5:**
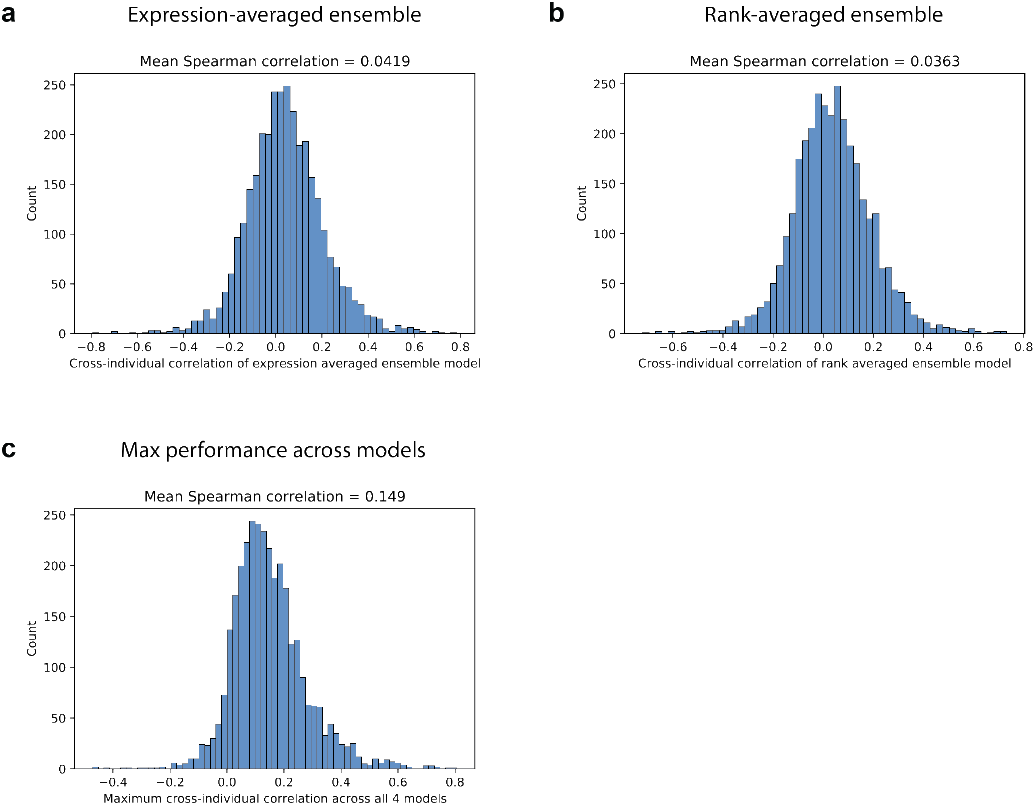
Performance of ensembled models. Predictions from the four models (Enformer, Basenji2, ExPecto, Xpresso) are ensembled per gene per individual by averaging (a) the z-scored expression predictions or (b) the cross-individual expression ranks from each model. We then compute the cross-individual correlations for these ensemble models by comparing the ensemble predictions to the measured Geuvadis expression levels for each gene. Additionally, in (c), we show the distribution of the maximum cross-individual performance (Spearman R) per gene across the four deep learning models (from Fig. S3).

**Fig. S6:**
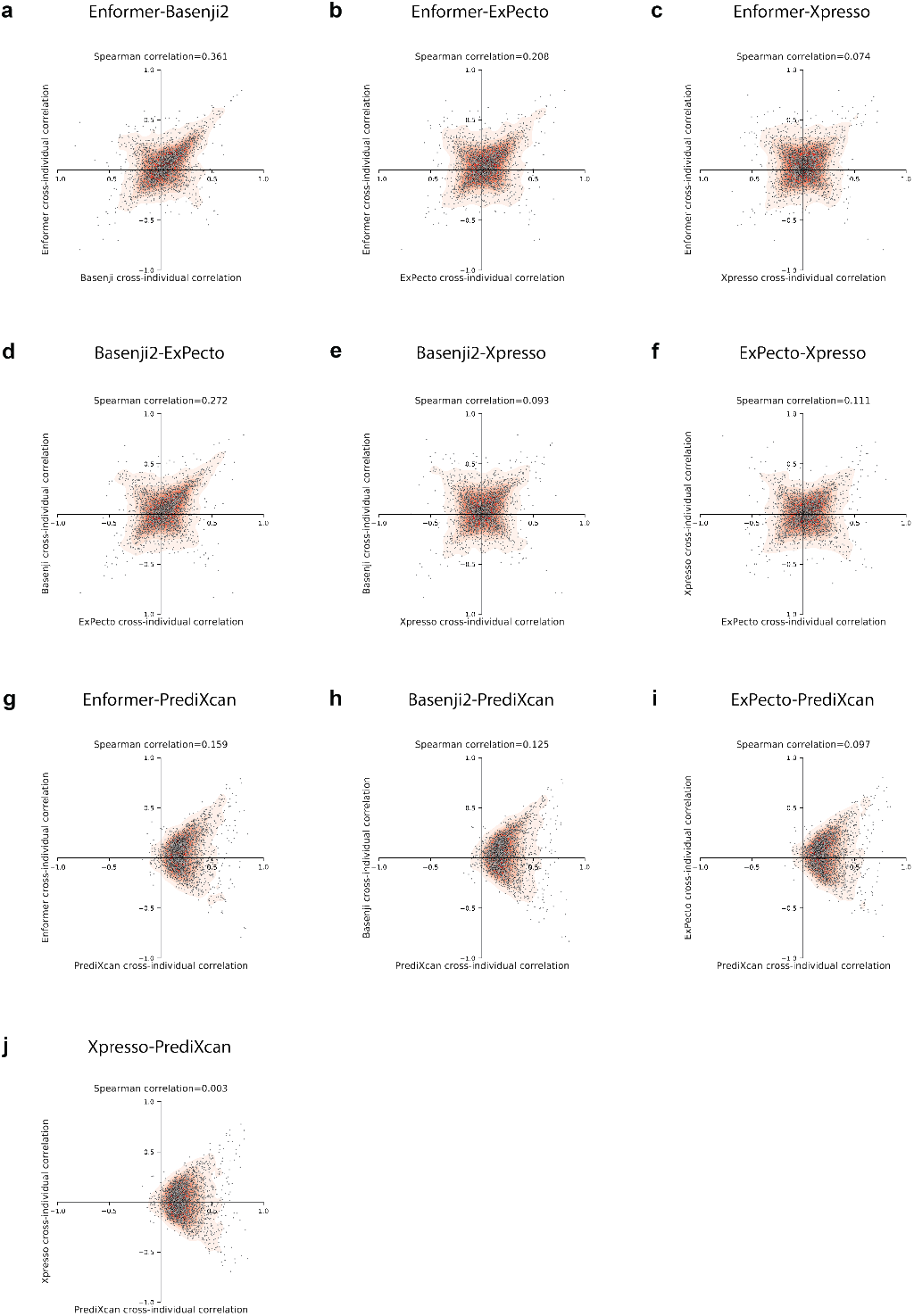
Pairwise model comparisons of cross-individual correlation. Comparison of cross-individual Spearman correlations between each pair of models. The scatter plot displays, for each gene, the performance achieved by both models, and a kernel density estimate of the scatterplot is overlayed on top. Note the increased density of genes along the *y* = *x* and *y* = *−x* axes.

**Fig. S7:**
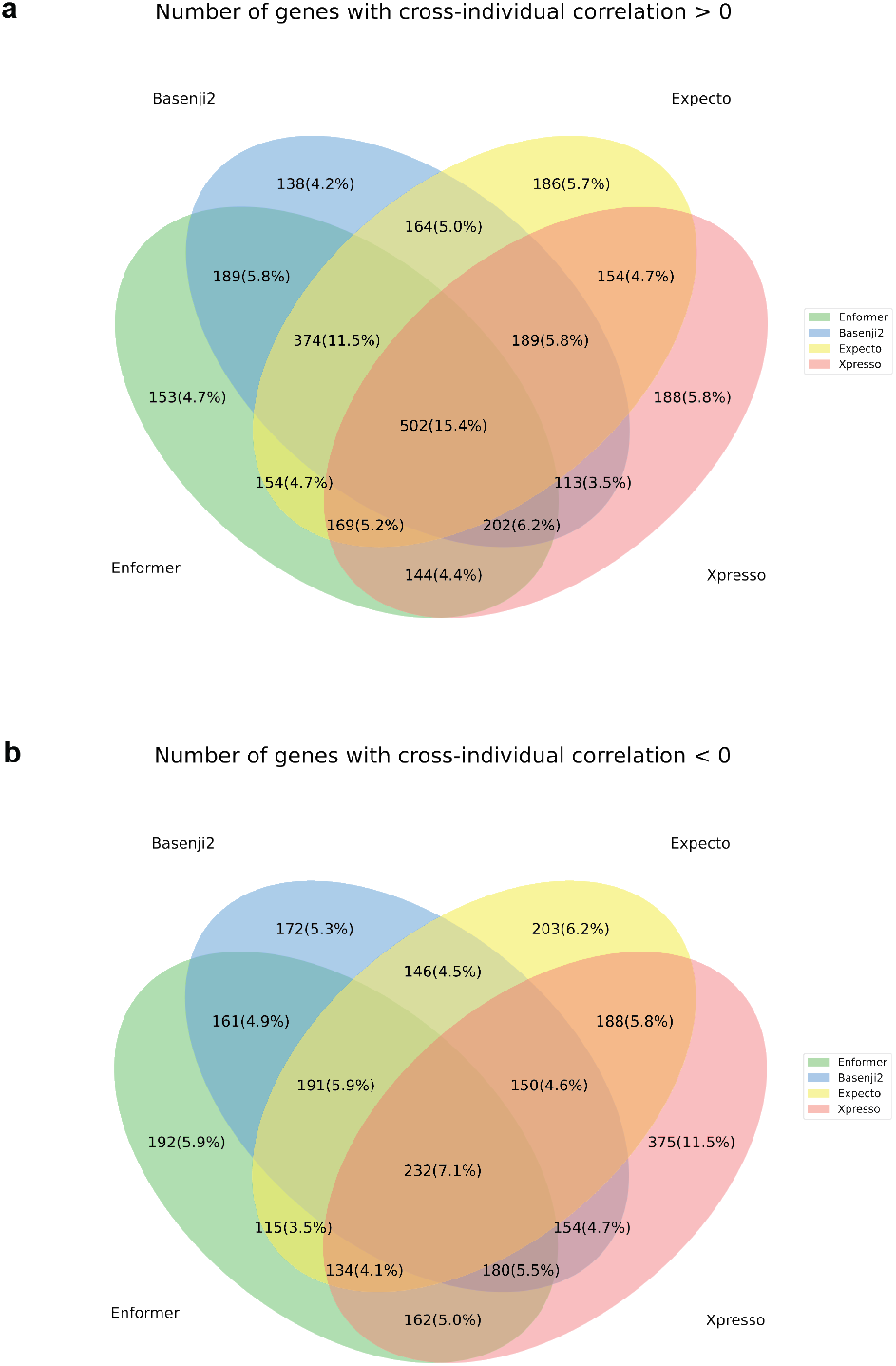
Consistency among methods. We display the overlap in (a) genes with a positive cross-individual correlation and (b) genes with a negative cross-individual correlation across the four evaluated methods. Percentages are out of the 3259 genes evaluated throughout this paper.

**Fig. S8:**
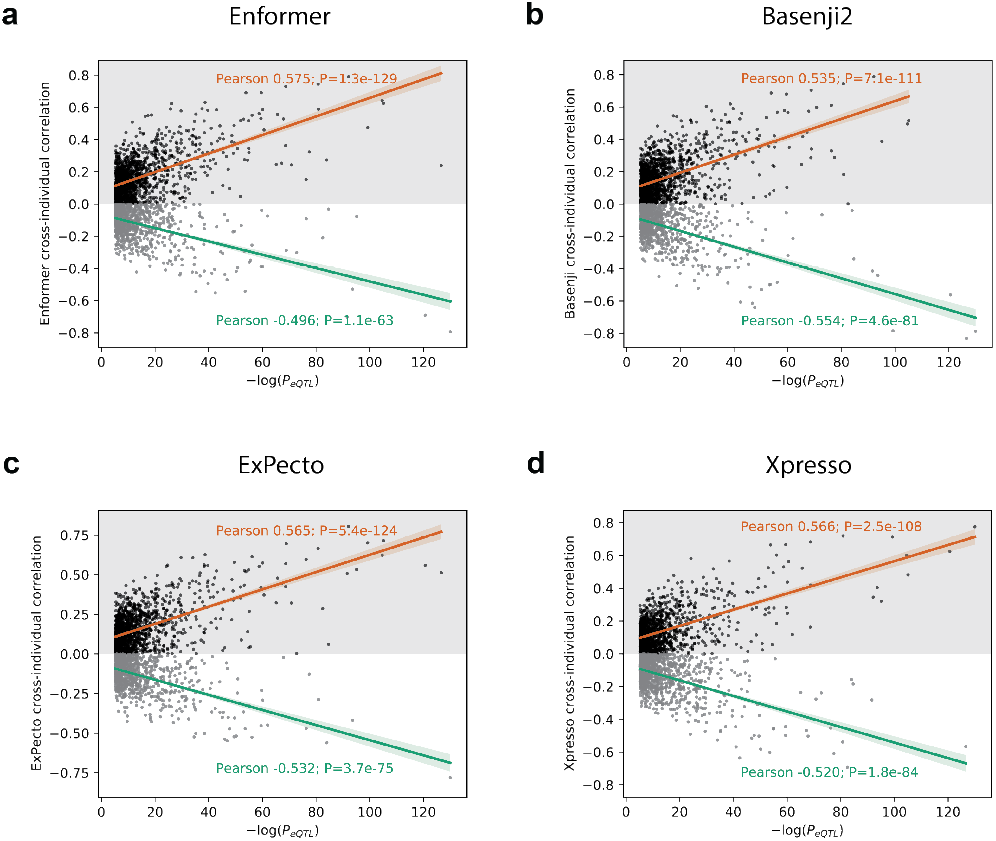
Cross-individual correlation vs. top eQTL p-value for all models. Cross-individual correlations for each model compared to the p-value of the most significant Geuvadis eQTL in each gene. For all models, we separately fit a line for genes with positive and negative cross-individual correlations. We report the Pearson’s *r* and p-value (two-sided Pearson correlation test) of each relationship.

**Fig. S9:**
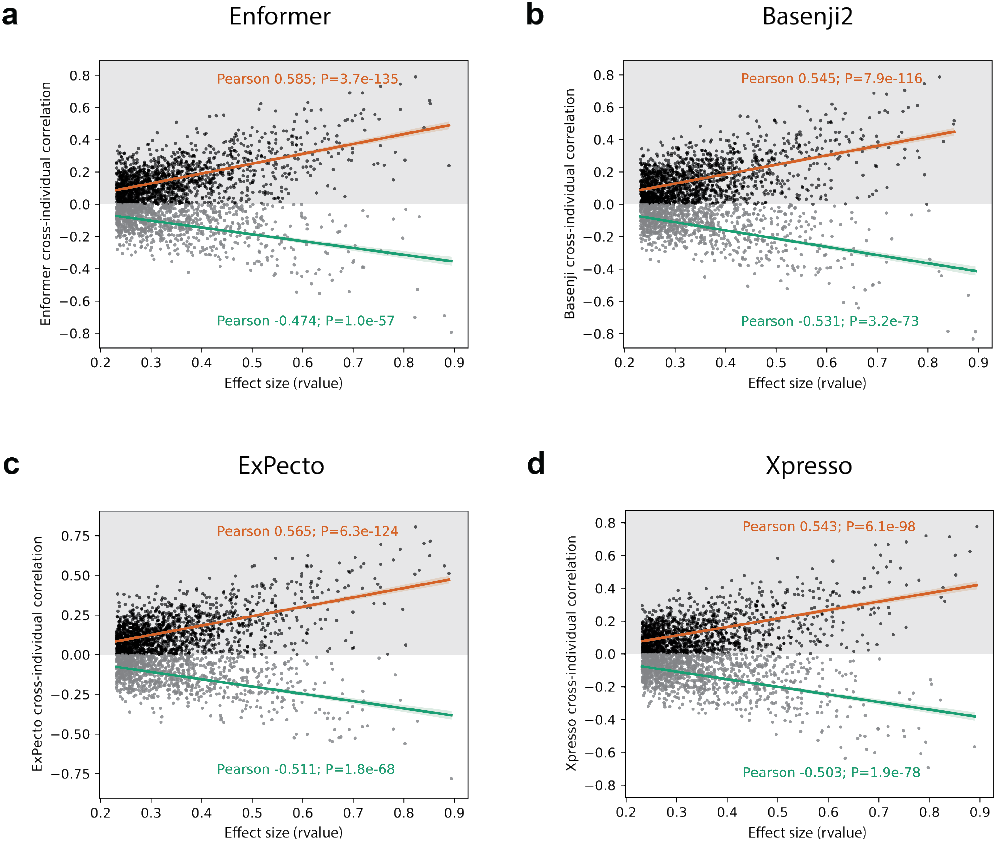
Cross-individual correlation vs. top eQTL effect size for all models. Cross-individual correlations for each model compared to the absolute value of the effect size of the most significant Geuvadis eQTL in each gene. For all models, we separately fit a line for genes with positive and negative cross-individual correlations. We report the Pearson’s *r* and p-value (two-sided Pearson correlation test) of each relationship.

**Fig. S10:**
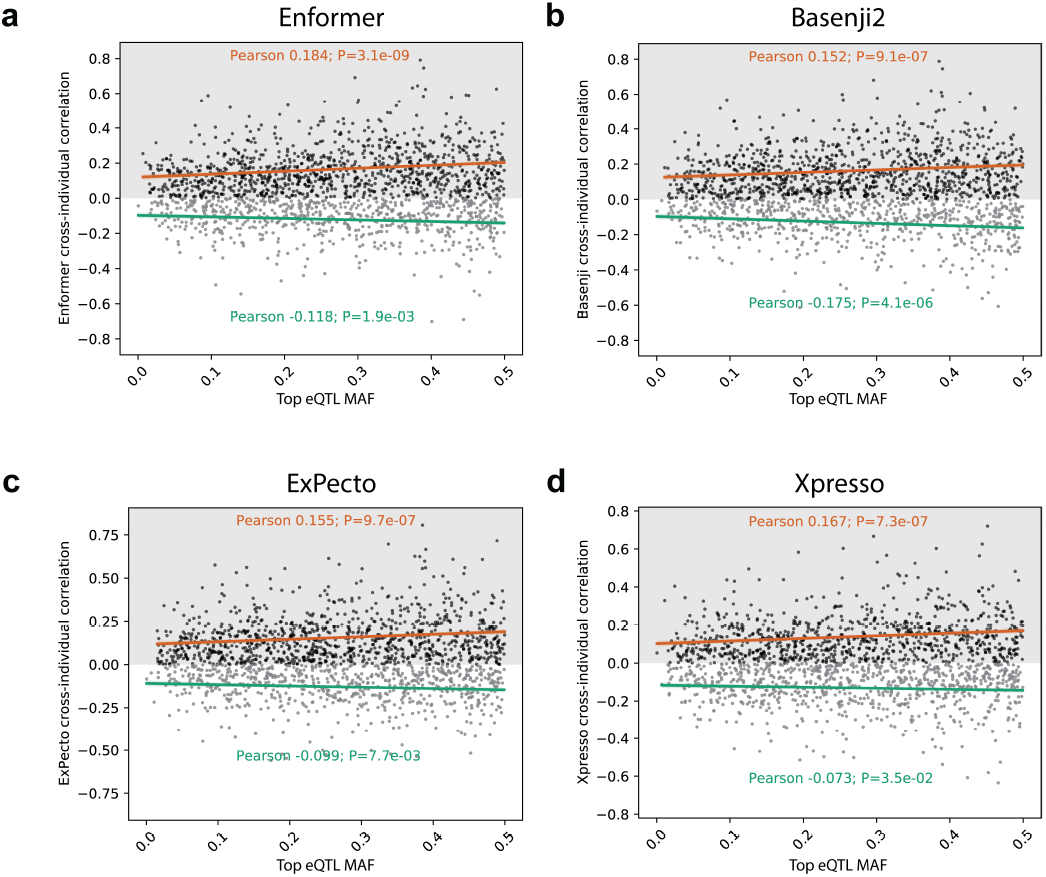
Cross-individual correlation vs. top eQTL allele frequency. Cross-individual correlations for each model compared to the global minor allele frequency (from Ensembl biomaRt) of each gene’s most significant Geuvadis eQTL. For all models, we separately fit a line for genes with positive and negative cross-individual correlations. We report the Pearson’s *r* and p-value (two-sided Pearson correlation test) of each relationship.

**Fig. S11:**
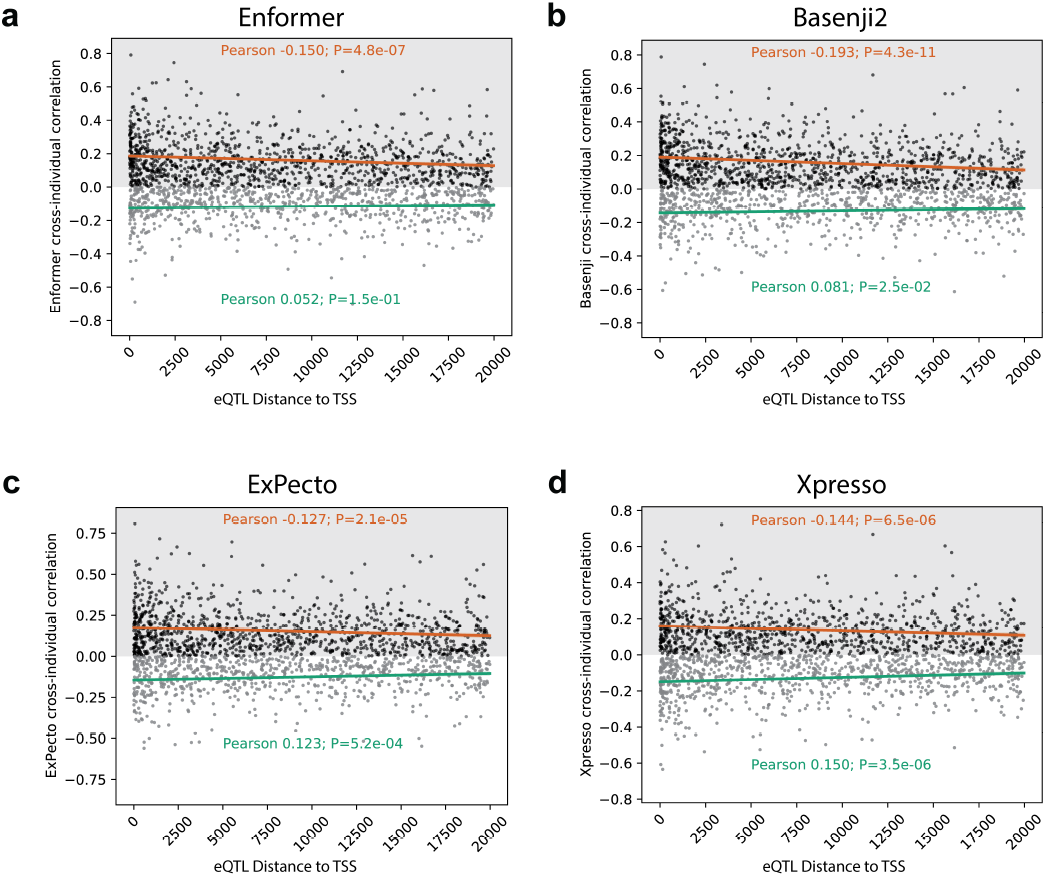
Cross-individual correlation vs. top eQTL distance to TSS. Cross-individual correlations for each model compared to the distance between each gene’s most significant Geuvadis eQTL and its TSS. For all models, we separately fit a line for genes with positive and negative cross-individual correlations. We report the Pearson’s *r* and p-value (two-sided Pearson correlation test) of each relationship.

**Fig. S12:**
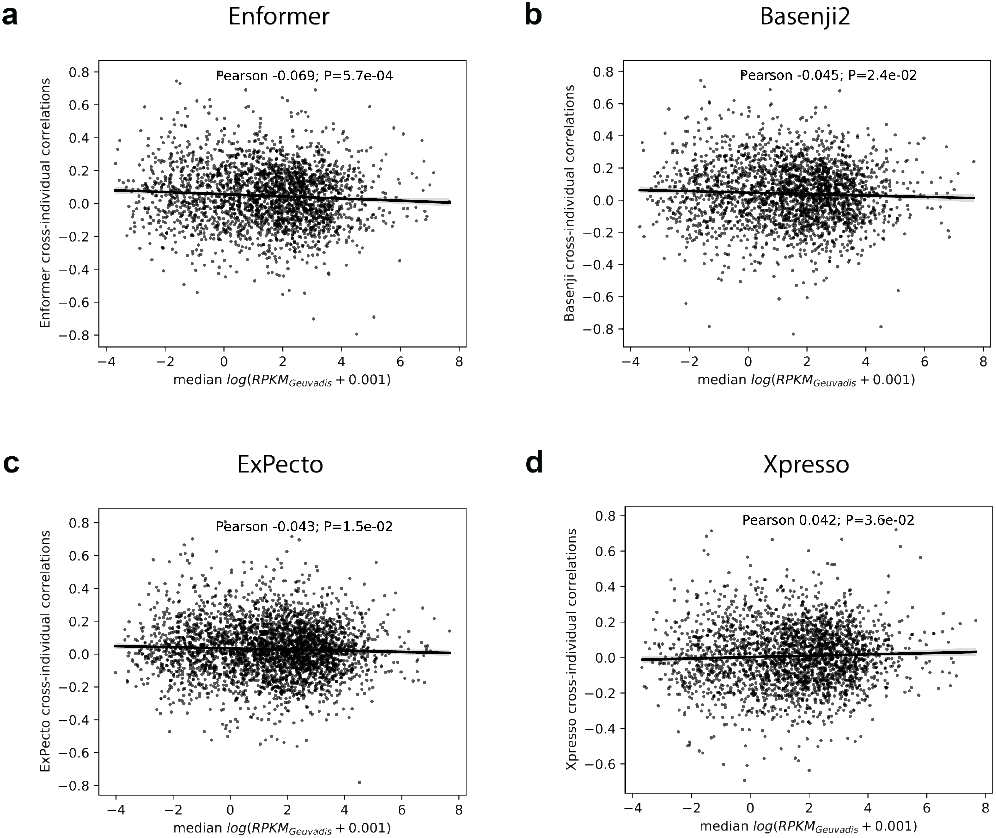
Cross-individual correlation vs. median gene expression for all models. For each model, we display the cross-individual correlations compared to the median Geuvadis gene expression level for each gene.

**Fig. S13:**
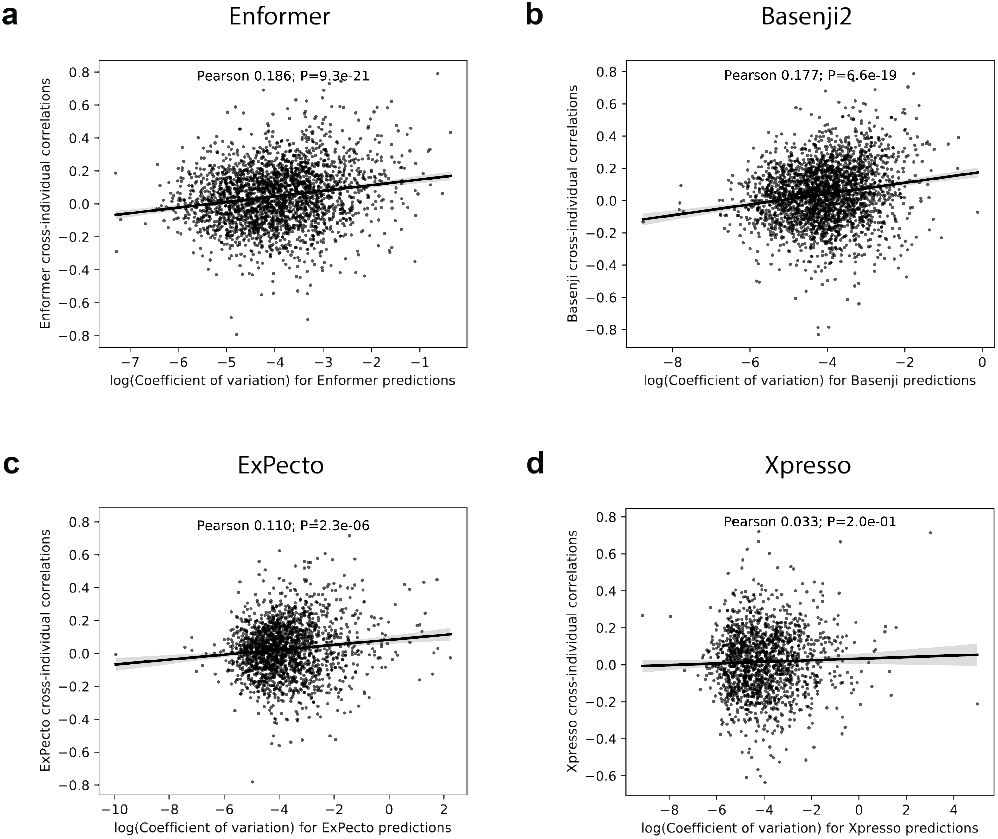
Cross-individual correlation vs. predicted expression dispersion for all models. For each model, we compare the cross-individual correlations to the log coefficient of variation 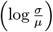,a measure of dispersion, in model predictions for each gene.

**Fig. S14:**
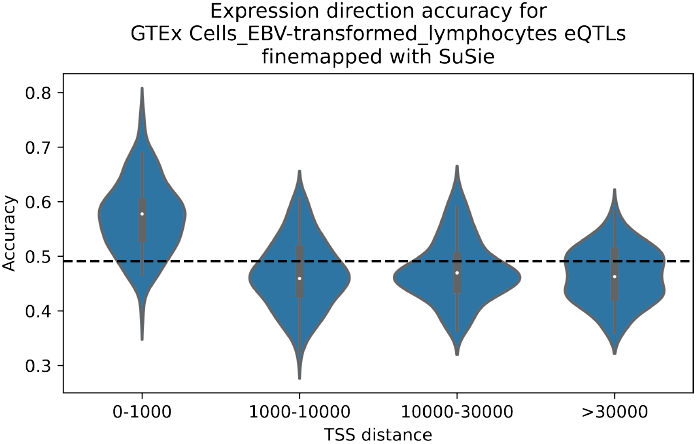
Enformer accuracy on predicting expression effect direction for fine-mapped eQTLs. Enformer expression direction prediction accuracy for GTEx EBV-transformed lymphocyte cell eQTLs fine-mapped using SuSiE. Enformer predictions from the GM12878 CAGE track were used to predict whether the minor allele for each fine-mapped eQTL increases or decreases gene expression. Accuracy on this task is shown stratified across four bins of variant distance to TSS, together with the mean accuracy over all distances (dashed line). Violin plots show the accuracy distribution of 100 bootstrap samples from the full set of variants.

**Fig. S15:**
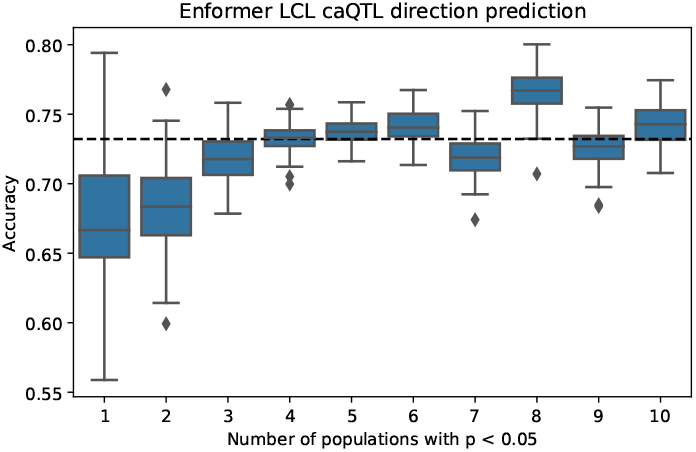
Enformer accuracy on predicting effect direction of LCL caQTLs. Enformer’s performance at classifying alleles as more or less accessible than the reference, using variants from the Tehranchi *et al*. dataset [21] in which chromatin accessibility quantitative trait loci (caQTLs) in LCLs were called from ATAC-seq data. We stratify performance by the number of populations (out of ten: four African, four European, one African-American, and one Han Chinese) in which the caQTL is identified as significant (*p <* 0.05). The dashed black line indicates average performance across all bins. Boxplots show the accuracy distribution over 100 bootstrap samples of the data. Note that these accuracies should not be compared directly with those for finemapped eQTLs reported in Fig. S14 because of substantial differences in the two methods for calling QTLs.

